# Context dependent multisensory integration: Mechanosensation depends on luminance for robust performance

**DOI:** 10.1101/2023.09.29.560224

**Authors:** Varun P Sharma, Simon Sponberg

## Abstract

Multisensory, goal-directed behaviors are ubiquitous and essential in animals across taxa. Individual sensory modalities may be significantly tuned by environmental context, and yet they must continue to combine and produce multisensory behaviors. This necessitates either behavioral robustness or goal-relevant adaptation in sensorimotor systems. In the case of hover-feeding hawkmoths of the species *Manduca sexta*, proboscis mechanosensation and vision are known to combine linearly in tracking of moving flowers. This tracking behavior is affected by light-levels, and visual compensation is necessary for robust flower-tracking. Whether the mechanosensory response also adjusts to light-level, in order to produce linear combination across luminance contexts, is not known. We applied control theoretic analyses to study integration of mechanosensory and visual information in flower tracking hawkmoths in the context of high luminance. This was performed using a two-part robotic flower system. Under the conditions of high luminance, we verified the linear combination of mechanosensory and visual systems, and the complementary modulation of these systems. We show that both systems tune their frequency response with changing environmental light level,while keeping their combined error low at relevant frequencies. Thus, the animal adapts to changing conditions, across modalities, even when only one of the modalities (vision) is directly affected. This cross-talk between sensorimotor systems is constrained, and possibly driven, by performance requirements.

## Introduction

Animal behavior often requires the integration of multiple sensory cues in a task-specific manner. This necessitates sensory-to-motor transformations which may simultaneously depend on the context and activity of other sensory modalities. Multisensory behaviors become especially challenging when a task must be performed across contexts (such as day and night) because the context can affect sensory processing.If the organism maintains performance across contexts which necessarily change one modality, then its other modalities must either be robust to these changes or also adjust when the context changes. Many organisms modulate one sensory modality when the reliability or salience of another modality changes. But how do organisms that behaviorally integrate multiple cues with different weights across frequencies tune the overall dynamic frequency as contexts change?

Control theory and system identification can help us identify how multiple modalities have changed across their dynamic range and whether their integration is robust or adaptive. Control theoretic experiments have revealed principles of multisensory integration, especially in reference-tracking tasks, such as cockroach wall-following (***Cowan et al. (2006***)), knifefish refuge-tracking (***Sutton et al. (2016***)), blowfly gaze stabilization (***Hengstenberg (1988***),***Hengstenberg et al. (1986***)), and hawkmoth flower-tracking (***Sprayberry and Daniel (2007***)). The electric knifefish, *eigenmannia virescens*, combines visual and electrosensory cues to track refuges, linearly superimposing the two modalities but adjusting their weights based on salience (***Sutton et al. (2016***)). Visual and tactile integration studies in human posture control (***Oie et al. (2002***)) and height-estimation (***Ernst and Banks (2002***)) have shown evidence for re-weighting of modalities depending on their salience. These are statistically-optimal strategies for the specific cues considered, which therefore have amplitude-dependent gains. However, they do not examine the strategy for combining cues across their dynamic range of frequencies. While the more general case of nonlinear integration is important (***Huston and Krapp (2009***),***Cellini and Mongeau (2022***)). in some animals the combination of cues is effectively linear, such as for hawkmoth flower-tracking (***Roth et al. (2016***)) and in *Drosophila* wing motor output (***Rauscher and Fox (2021***)). The respective contributions to behavior of the different sensory modalities in these studies was investigated in a single behavioral and environmental context. However, sensory systems are tuned to their environment and have characteristic timescales ***Roth et al. (2016***); ***Dahake et al. (2018***). Two strategies are possible for maintaining this performance across contexts. A robust combination means that one modality does not necessarily have to change in conjuction with the other to maintain behavioral performance across the dynamic range. An adaptive strategy means that there must be concomitant changes in one modality when the other adjusts for context.

*Manduca sexta* is a crepuscular hawkmoth, capable of hover feeding in mid-air while tracking flower movement. Flower-tracking behavior involves closed-loop flight control, with frequency-response dynamics that depend on sensory modalities. Using dummy flowers and controlled sensory stimuli, we know that these flower-tracking dynamics are linear (***Sprayberry and Daniel (2007***),***Sponberg et al. (2015***), ***Dahake et al. (2018***),***Stöckl et al. (2017***)), and this tracking is performed using visual and proboscis mechanosensory information. Visual and antennal mechanosensory contributions to flower-tracking are widely separated in frequencies (***Dahake et al. (2018***)). Proboscis mechanosensation, however, has significant overlap with visual flower-tracking response. ***Roth et al. (2016***) used robotic flowers to infer a template for multisensory integration of visual and proboscis mechanosensory inputs, as a linear combination of the two systems. They also found a frequency dependent separation of the role of each sensory system, with the visual system sensitive to relatively high frequencies. These results were tested for a single, low luminance level. Decrease in luminance results in an increase in time-lag of the moth’s tracking of a moving flower (***Sponberg et al. (2015***)). Therefore, a change in the visual context has an effect on the moth’s behavior, while also directly affecting at least one of the two sensory systems used in this behavior. Animal locomotor behavior is often performed in changing environmental conditions, such as light-level, terrain roughness or wind-speed, which are very different contexts. Changing context can result in a change in the gain (magnitude) or the sensitivity of a sensory response. Context dependence of neural systems has been demonstrated multiple times, such as by ***Maimon et al. (2010***), ***Olberg (1983***), ***Sponberg et al. (2015***),***Stöckl et al. (2016***) and ***Theobald et al. (2010***). In ***Maimon et al. (2010***), changing the context, sometime termed the behavioral state, of a fly from stationary or flapping changed the gain of motion-processing neurons. In addition to internal state, informational cues from the environment, such as different odors or light-levels, help determine the context for a behavior and change the tuning of sensory systems. In moths, this is seen in the gating of flip-flop activity by pheromones to drive cast-and-surge behavior (***Olberg (1983***)), and luminance-dependent adaptations for flower-tracking and optomotor correction (***Sponberg et al. (2015***),***Stöckl et al. (2016***),***Theobald et al. (2010***)).

It is not surprising that the gain and phase response of the visual system depends on background. Here, we test whether overall luminance (i.e. the changing context) also affects the hawkmoths’ mechanosensory response to a moving flower separately from its impacts on vision and how the two cues are combined in these different settings. To maintain tracking performance across the range of frequencies at which the moth track natural, wind-blown flowers ***Sponberg et al. (2015***), the animal may transform mechanosensory cues robustly, in which case we would expect to see the gain response of the mechanosensory pathway remain constant but to not significantly degrade performance. Alternatively, the mechanosensory system itself may also be luminancedependent in a way that it adapts sensorimotor control to maintain performance in the new context. In either case, both visual and mechanosensory systems depend on luminance, their linear combination across light-levels may persist or may switch to a non-linear integration. Finally, these changes may degrade behavioral performance, or the moths may track flowers equally well or better despite the frequency-dependent modulation of one or both sensory modalities.

## Results

In multisensory behaviors, probing a system’s response to conflicting sensory information can separate the sensorimotor transform for each cue and show how they combine. We used sensory-conflict and coherent-motion stimuli, as described in Roth et al (***Roth et al. (2016***))[see Methods and Materials], but now across contexts. Briefly, our setup consists of a two-part robotic flower, with a facade (V) and a nectary (M) for visual and proboscis mechanosensory inputs, respectively (Figure 1a). In order to test for context dependence, we performed these experiments at the flower illumination value of 300 lux (early dusk) instead of the previously used value of 0.3 lux (starlit night). Based on the observations at 0.3 lux, we hypothesized that at 300 lux the two modalities combine linearly, as per the control diagram in Figure 1a. For system-identification, we provided sum-of-sines stimuli (see Methods and Materials) in M-moving (facade stationary), V-moving (nectary stationary) and coherent motion (both moving together) cases (Figure 1b). Moth trajectories in response to these motion stimuli were recorded. In the frequency domain, like in previous studies, we found that Fourier amplitude peaks are present selectively at each of the driving-frequencies, up to at least 3 Hz for all 12 individuals and all 3 stimulus conditions (Figure 1b). Using combinations of these three experiments we can isolate the dynamics of the mechanosensory system separate from the visual system across luminance conditions.

**Figure 1.**
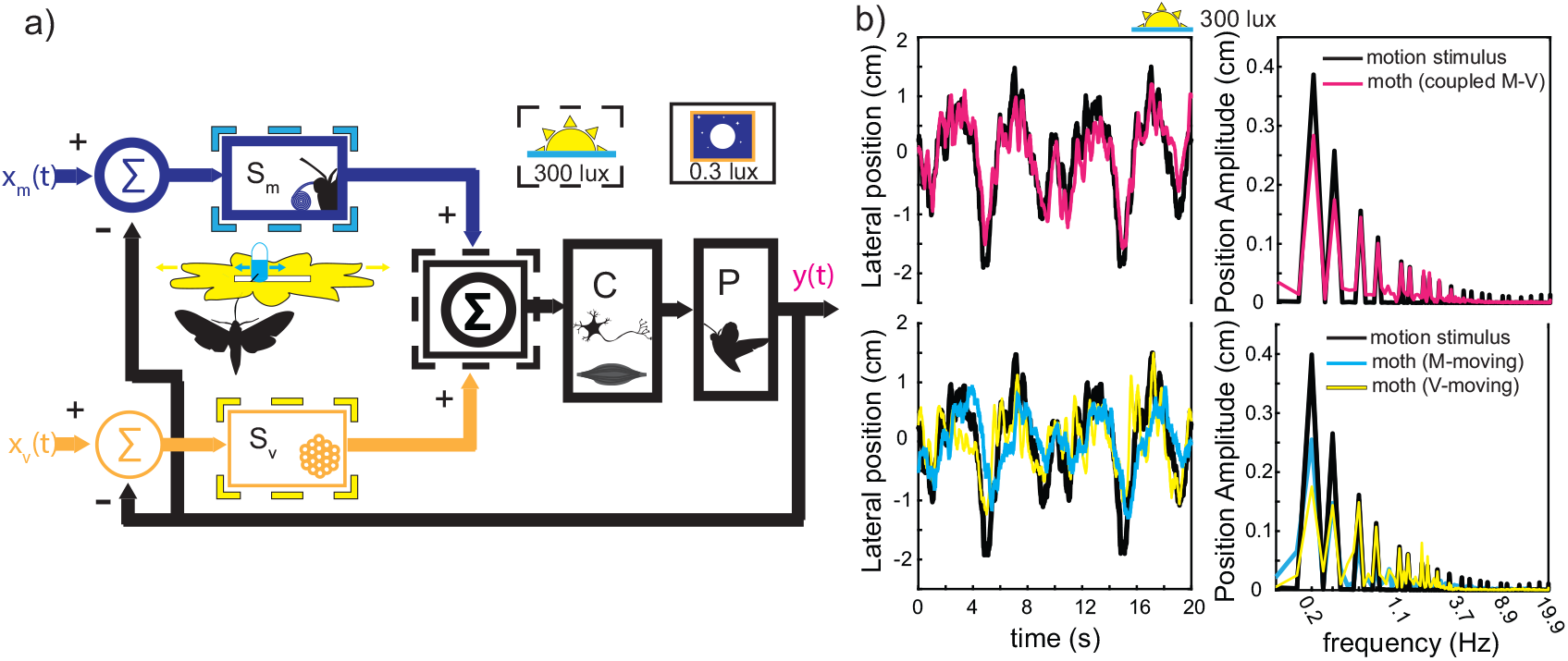
(a) Feedback control diagram for multisensory integration. Under conditions of high-luminance (300 lux, lighter shades, dashed boxes) and low-luminance (0.3 lux, darker shades), the visual (yellow-orange) and mechanosensory (blue-cyan) systems are summed linearly. Mechanosensory and visual position stimuli (r_*m*_ and r_*v*_) result in moth positional response (y, purple-pink). The sensory (S_*m*_ and S_*v*_), control (C) and plant (P) blocks correspond to sensory, neural and biomechanical subsystems. (b) Time-domain and frequency-domain flower-tracking at high (300 lux) luminance. The coherent MV response (pink), as well as M-moving (cyan) and V-moving (yellow) responses, to motion stimuli (black) show a reliance on both sensory systems. Responses have significant peaks at several driving-frequencies

### Mechanosensory and Visual Systems Combine Linearly at High Luminance

We first test whether the visual and mechanosensory cues are still combined linearly across contexts (luminance levels). Under each stimulus condition (V-moving, M-moving, and MV-moving), we quantify the moth’s response to flower motion at each frequency in stimulus using the gain and phase delay in the response. Gain and phase define points in the complex plane as polar coordinates. Frequency responses provide a simple way to test the hypothesis because if the two cues combine linearly then simply adding (in the complex plane) the response to the V-moving and M-moving experiments should exactly predict the MV-moving experiments. This is not trivial because in the V-moving and M-moving cases the other cue (the nectary or the facade of the flower, respectively) is stationary and so the moth must balance conflicting cues in each case. In order to check consistency of linearity, we first measured the M-moving and V-moving closed-loop transfer-functions (Hm and Hv), defined by the gain and phase response in the M-moving and V-moving conditions. These tell us how the moths respond to sensory conflict, in closed-loop. The mechanosensory and visual closed-loop transfer functions are given by:

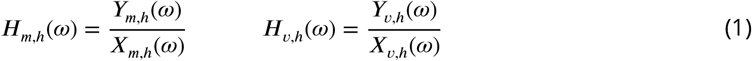

Here, Y_m,h_(*ω*) and X_m,h_(*ω*) are the measured lateral positions of the moth and flower respectively, in the M-moving case, while Y_v,h_(*ω*) and X_v,h_(*ω*) are for the V-moving case, all in the frequency-domain. The subscript ‘h’ denotes high luminance (300 lux). We compare these high-luminance transfer function measurements with their low-luminance counterparts. If one or both of these transfer functions change with luminance, then the system has luminance-dependent adaptation.

We find that the closed-loop response (H_m_ and H_v_) for both M-moving and V-moving stimuli, changes with luminance. For mechanical-moving stimuli, the gain is significantly higher (p=0.95, t-test on gain and phase at each frequency) in the low luminance condition (Figure 2a) than for high luminance. The opposite is true for visual-moving stimuli (Figure 2b), where the gain is higher for high luminance, over the range of ecologically relevant frequencies (see ***Sponberg et al. (2015***) for natural frequencies of flowers). Despite these differences, the M-moving response has a low-pass characteristic for both luminance conditions, while the V-moving response is more akin to band-pass (i.e. has a distinct peak in gain) for both luminance conditions. The V-moving peak-gain frequency shifts from 2.9Hz (at 0.3 lux) to 1.9Hz (at 300 lux). Therefore, the moth’s multisensory flower-tracking system has quantitative luminance-dependent adaptation, while keeping the qualitative shapes of the transforms. Given this adaptation, it is not trivially obvious that H_m_ and H_v_ will continue to combine linearly.

**Figure 2.**
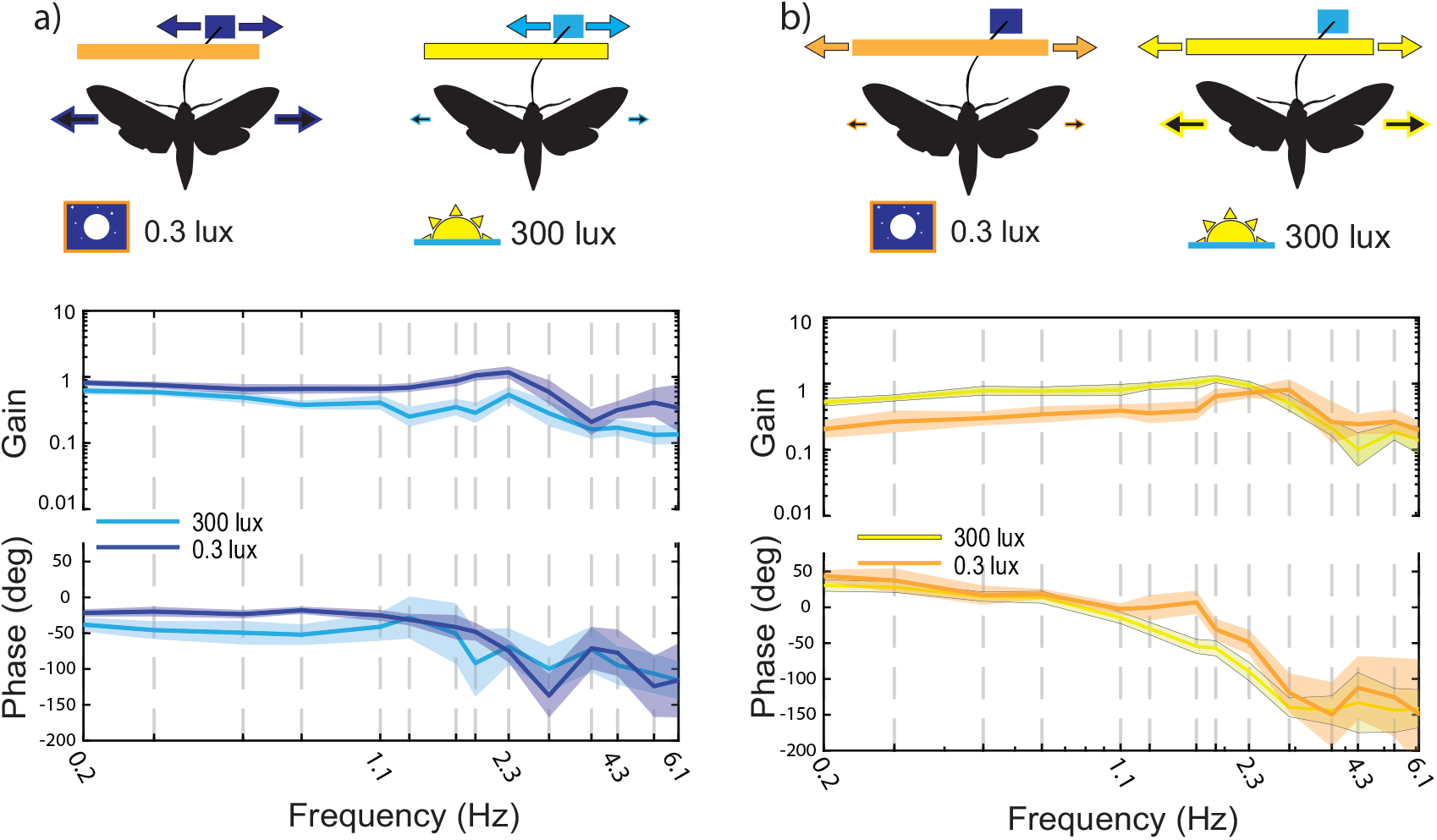
Closed-loop Mechanosensory (a) and Visual (b) motion frequency response functions. Gain and phase of the M-moving (blue-cyan) and V-moving (yellow-orange) transfer functions in (a) and (b) respectively, show luminance-dependence. The lighter shades correspond to 300 lux and the darker shades correspond to 0.3 lux. Mean and 95% confidence intervals are shown, calculated as described in Methods and Materials. Illustrations above the plots show top-views of the moth, nectary and facade, with arrows indicating motion

To test linear combination, in addition to sensory-conflict, we also measured the response of moths to coherent motion of stimuli (MV-moving):

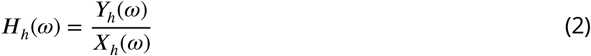

Here, Y_h_(*ω*) and X_h_(*ω*) are the frequency domain representations of the moth’s and coherent-stimulus’ positions, respectively, as a function of frequency (*ω*). At low luminance, Roth et al ***Roth et al. (2016***) found that mechanosensory and visual contributions sum linearly, as in Figure 1a (control diagram) and Figure 3a (measured and predicted bode plots). To ascertain whether linear summation of modalities holds true at high luminance (300 lux) as well, we calculated the mean linear-sum prediction:

**Figure 3.**
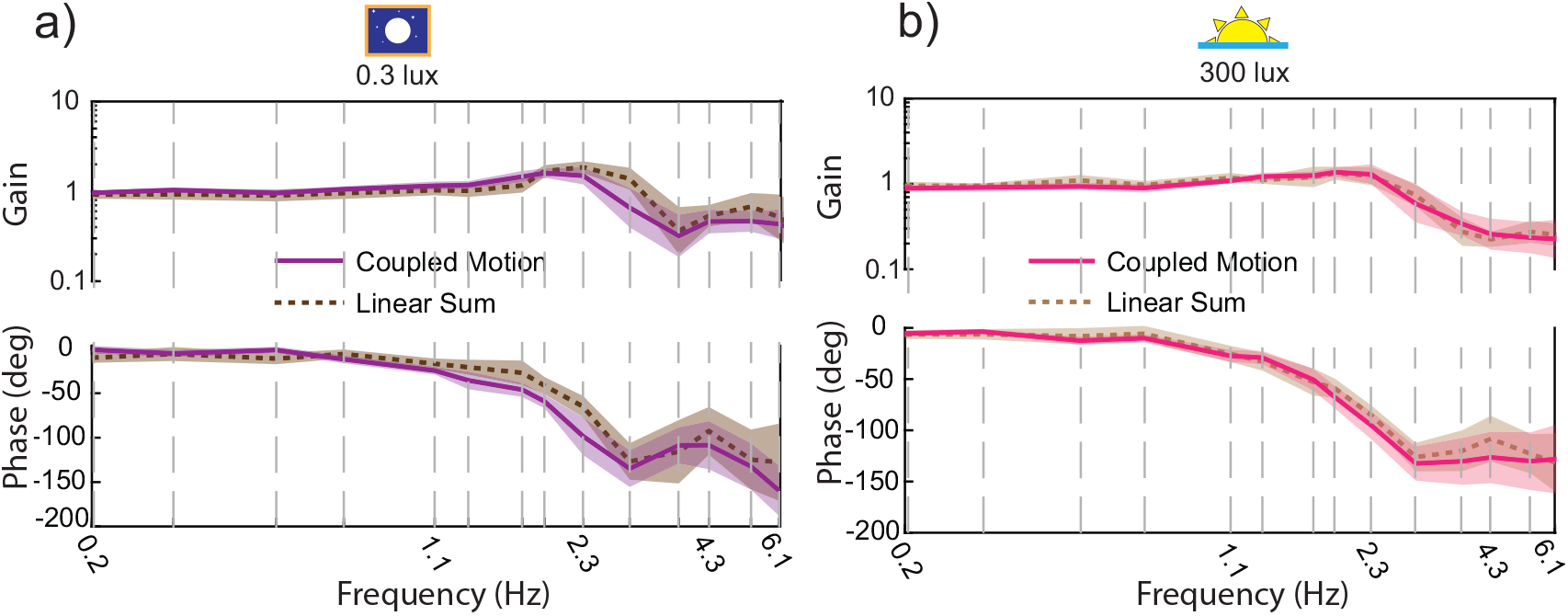
Linear Sum Validity Across Luminance Contexts: Gain and phase plots for predicted (brown) and experimental (violet-pink) coherent-tracking transfer function at (a) 0.3 lux (*H*_*l*_) and (b) 300 lux (*H*_*h*_). Mean and 95% confidence intervals of the mean are shown, as described in Methods and Materials

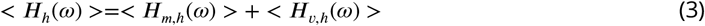

On comparing this with the mean transfer function for coherent motion (H_h_(*ω*)), we find that the two are not significantly different in both gain and phase (Figure 3b, 95% confidence intervals of mean, at each frequency). Next, we evaluated the linear-sum prediction individually for each moth, which states that H_m_ and H_v_ sum to give H, the coherent tracking response. For all individuals considered, the linear-sum prediction matches the experimental coherent motion result over the range of ecologically relevant frequencies. Hence, frequency-dependent re-weighting of mechanical and visual systems maintains the linearity of their integration, at every frequency. The summation-template for the eyes and proboscis, described by ***Roth et al. (2016***) (Figure 1a), persists at illumination levels 1000-fold higher than the one used in their study.

### Mechanosensory and visual systems are separately modulated by luminance

The observation that H_v_ and H_m_ change with luminance does not tell us whether both visual and mechanosensory systems are individually modified. This is due to the nature of closed-loop measurements, since H_m_ is affected by the visual sense as well due to sensory conflict. In order to test if the observed luminance-dependent adaptation was due to shifts in only the visual system, or involved changes in both vision and mechanosensation, we must mathematically “open” the closed loop behavior. We do this by combining the behavior results from the M-moving, V-moving and coherent-motion conditions and factoring out the changes due only to the mechanosensory and only to the visual systems, which we define as the open-loop mechanosensory transform, G_m_, and the open-loop visual transform, G_v_, respectively (Figure 4). Opening the loop to evaluate G_m_ and G_v_ requires comparing across stimulus conditions. These transforms tell us how moths react to mechanosensory and visual errors, individually (Figure 4).

**Figure 4.**
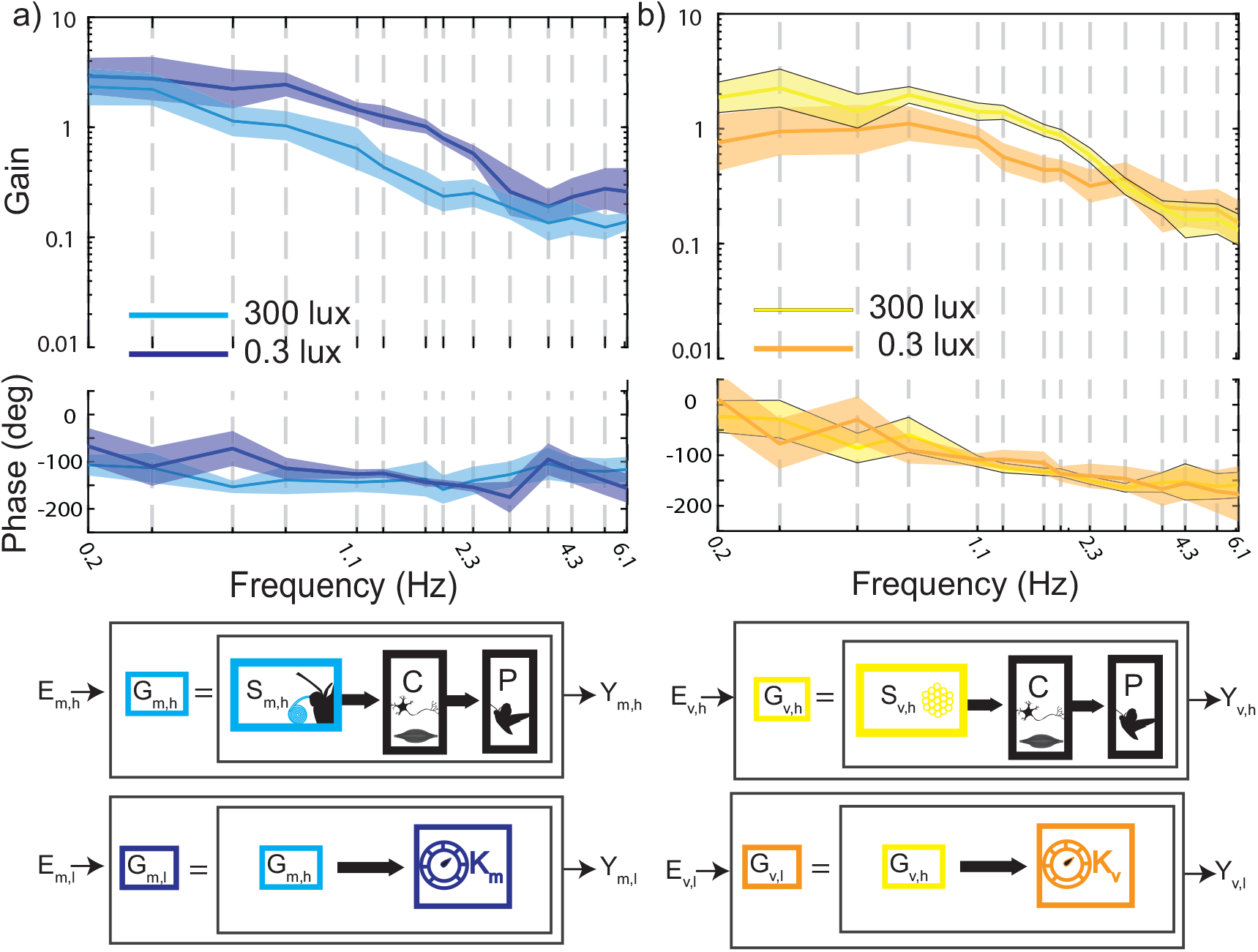
Both visual and mechanosensory open-loop systems change with luminance: Frequency response functions for (a) mechanosensory (*G*_*m,l*_: blue, and *G*_*m,h*_: cyan) and (b) visual (*G*_*v,l*_: orange and *G*_*v,h*_: yellow) open-loop transfer functions. The high luminance (300 lux, lighter shades) open-loop transfer functions (mean and 95% confidence intervals) are calculated for each individual, as described in the text. The low-luminance (0.3 lux, darker shades) open-loop transfer functions (mean and 95% confidence intervals) are calculated through permutation, as described in the text. The open-loop transfer functions transform the stimulus-error (E_*m*_ and E_*v*_) into moth positional response (Y_*m*_ and Y_*v*_) by action of sensory, neural and biomechanical blocks (S, C and P). High to low luminance transformation is composed of frequency dependent gain changes (*K*_*m*_ and *K*_*v*_)

The (experimentally measured, Figure 2) closed-loop transfer functions H_m_ and H_v_ are related to the open-loop transfer functions (G_m_ and G_v_) by:

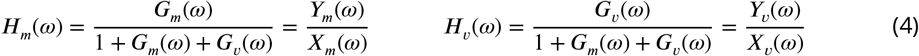

Thus, we can infer the open-loop transfer functions using the closed-loop transfer functions measured from conflict (Eq 4) and coherent (Eq 2 and 3) trials:

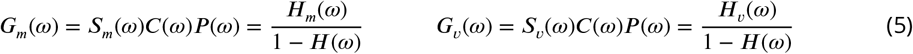

In the following equations, the subscript ‘h’ indicates that they pertain to the high luminance experiments (300 lux).

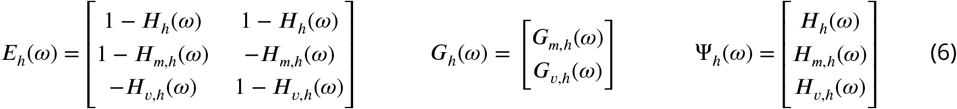

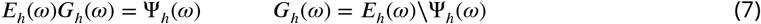

For each frequency (*ω*), and for every individual moth, we thus have an estimate of the open-loop mechanosensory (G_m_) and visual (G_v_) transfer functions (Eq. 7). Here, we have used MATLAB’s *mldivide* function (“\” operator). We performed a similar calculation on low luminance data (Methods and Materials), but with unpaired data across individuals

On comparing the estimates for open-loop mechanosensory transfer functions, *G*_*m,h*_ and *G*_*m,l*_ (Figure 4b), we find that mechanosensory gain is decreased for high luminance, and this decrease is most apparent between 0.5Hz and 2.9Hz. At 1.3Hz, the mechanosensory gain falls from 1 to around 0.4. The opposite trend is seen for the open-loop visual transfer functions *G*_*v,h*_ and *G*_*v,l*_. Visual gain is increased in the high luminance condition, while the phase is not significantly different between high and low luminance. The increase in visual gain is most apparent between 0.7Hz and 2.9Hz. The difference in mean gain for several frequencies up to 4.3Hz in both mechanosensory and visual cases is significant (t-test, p=0.95, evaluated at each frequency individually).

The mechanosensory open-loop gain changes with luminance. Therefore, we conclude that both mechanosensory and visual systems are individually dependent on light-level. The change in the two systems varies in a frequency-dependent manner. This is not an intuitively obvious result, since there is currently no known direct mechanism for modulation of proboscis mechanosensory output by luminance. Our result does not necessitate a change in the sensory response of the proboscis afferent neurons, but does predict a luminance-dependent change in the quantitative, motor response to proboscis stimulation. The shift, opposite to visual change, in these behavioral mechanosensory responses, may manifest itself in a number of ways, possibly through changes in neuromodulator profiles at multisensory-integration sites in the brain, and in the flight-control networks of the thorax.

### Visual and mechanosensory errors are changed, and coherent error is maintained constant across luminance levels

Since the mechanosensory and visual transfer functions depend on luminance, we next test whether flower-tracking performance is affected by these modifications. We defined complex tracking-errors for coherent and conflict stimuli, as was done by ***Roth et al. (2016***). The coherent and conflict errors are defined in the complex plane, where perfect tracking of the moving stimulus is gain 1 and phase 0 (at point 1+0j). Deviation from this perfect tracking point is the complex tracking error. Coherent motion error:

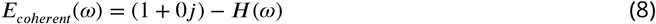

Error with respect to the moving stimulus:

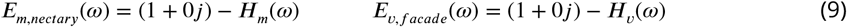

Error with respect to stationary stimulus:

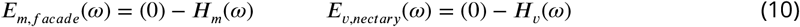

E_coherent_, E_m_ and E_v_ correspond to coherent (MV) motion, M-moving and V-moving conditions, respectively. The magnitude of coherent motion errors is interpreted as a performance metric. We found that coherent tracking error is not significantly different at 0.3 lux and at 300 lux (Figure 5a). The error magnitudes, when plotted for coherent motion (Figure 5a) show that error is small for low frequencies, greater than 1 at intermediate frequencies, and close to 1 for high frequencies (matching the trend seen in ***Sponberg et al. (2015***) and ***Roth et al. (2016***)). This means that the moth tracks slow-frequencies of flowers (less than 1Hz, similar to natural flower motion ***Sponberg et al. (2015***)) very well, at the expense of poor tracking at higher frequencies. Both high and low-luminance coherent-motion errors are very similar, with a characteristic overshoot around 2.3Hz. Hence, despite modulation of individual sensory responses, the combined response error is indistinguishable for luminance changes of 3 orders of magnitude.

**Figure 5.**
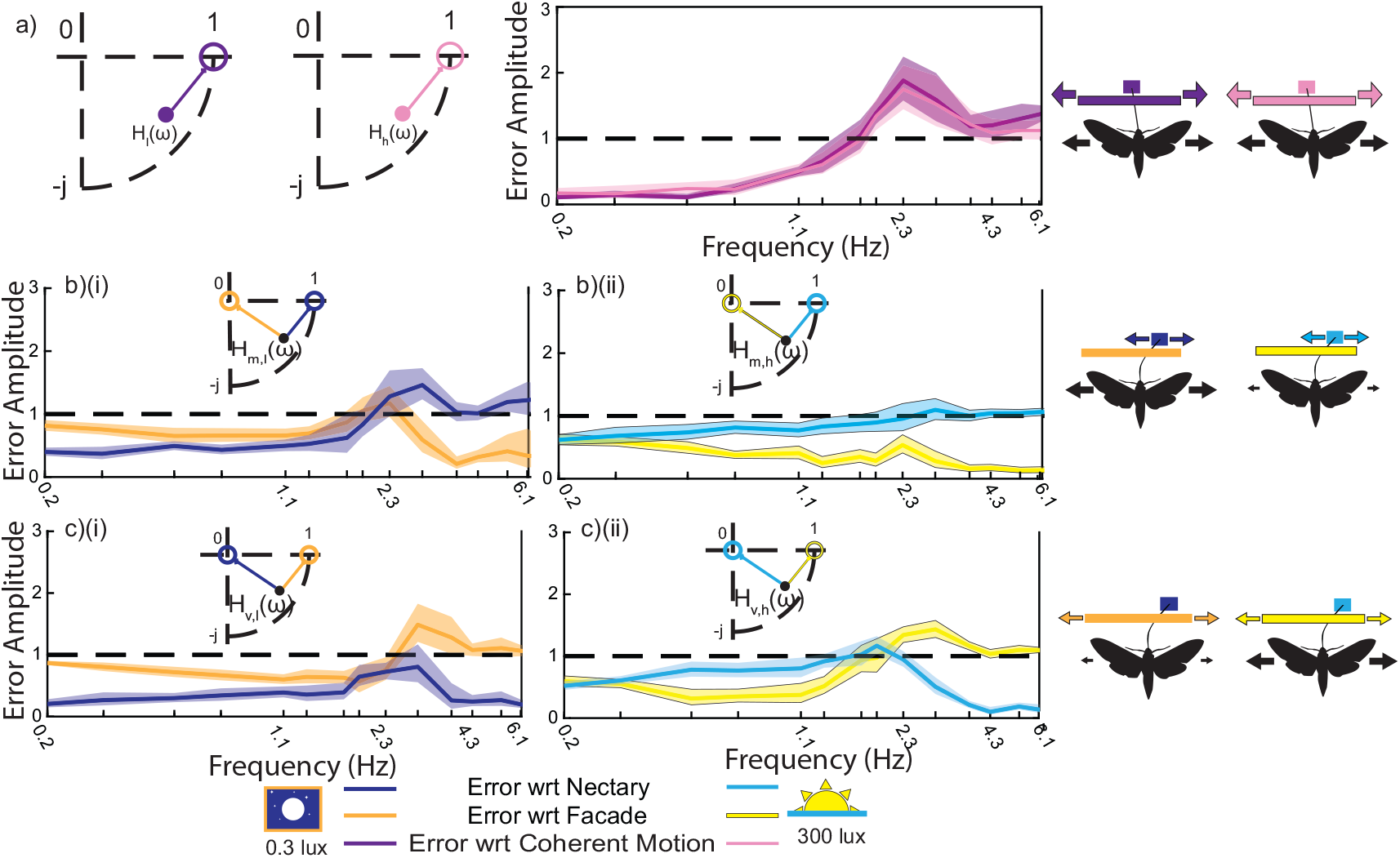
(a) Coherent tracking error is not significantly different for 0.3 lux and 300 lux. This is seen from the error magnitude for high and low luminance, violet-pink (b) Mechanical motion case: error magnitudes (with respect to facade and nectary) show speed-dependent differences in tracking error across contexts (c) Visual motion case: error magnitudes (with respect to facade and nectary) show speed-dependent differences in tracking error across contexts. Darker shades (i) correspond to 0.3 lux and lighter shades (ii) correspond to 300 lux. Means and 95% confidence intervals are plotted, as described in Materials and Methods

The individual error magnitudes of E_m_ and E_v_ can tell us which modality the hawkmoths weigh more strongly at different luminances. In the M-moving case, at 0.3 lux the mechanosensory error (E_m,nectary_) is lower than the visual error (E_m,facade_) up to 2.3Hz (Figure 5b). Meanwhile, the visual error is smaller at 300 lux, at all frequencies tested. In the V-moving case, a similar trend holds true (Figure 5c). Here, E_v,nectary_ is smaller than E_v,facade_ for most frequencies at 0.3 lux, while E_v,facade_ is smaller up to 1.9Hz at 300 lux. This relates to the greater reliance on the visual system at 300 lux, and the slower-speed tuning of the proboscis mechanosensory system. These observations are consistent with visual response having a higher weight at high luminance, and mechanosensory response at low luminance. Thus, despite a switch in the higher weighted modality, we find that coherent tracking performance is very similar across luminance conditions (Figure 5a), for the entire range of ecologically relevant speeds (less than 2Hz ***Sponberg et al. (2015***)). As the luminance decreases, the moth’s eyes and early visual system compensate using a spatial and temporal summation strategy (***Sponberg et al. (2015***), ***Stöckl et al. (2016***)), which may contribute to the increased error in response to facade motion. By increasing the relative weight of proboscis cues in this condition, coherent tracking performance is maintained.

### Luminance Dependent Delay is the Result of Multisensory Reweighting

Hawkmoths are known to effectively slow their brains to perform flower-tracking in low-luminance conditions (***Sponberg et al. (2015***),***Stöckl et al. (2016***), ***Stöckl et al. (2017***)). While a number of possible feedback-control models may fit this description, the result is consistent with a sensorimotor delay of 10ms for a luminance change from 300 lux to 0.3 lux. However this effective or emergent delay may results from multiple changes in the underlying sensory processing such as spatial and temporal summation at low light levels (***Warrant (1999***), ***Stöckl et al. (2016***), ***O’Carroll and Warrant (2011***)). Adding a delay term (*τ*=10ms) to the coherent open-loop response at high luminance, we compare it with the coherent closed-loop response at low luminance (Equation 13, where s=*jω*).

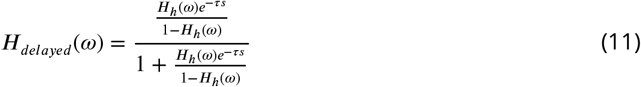

As in (***Sponberg et al. (2015***)), the delay has the effect of increasing the gain peak over the range of frequencies from (1.3 to 2.9 Hz), and the phase response rotates to drop by 3.6 degrees per unit frequency (deg/Hz), especially evident after 6.1Hz (Figure 6b). This prediction is not significantly different from the experimental data from low luminance trials at these frequencies. Unlike in (***Sponberg et al. (2015***)), the secondary peak present at 0.3 lux between 2.9Hz and 8.9Hz, disappears for 300 lux conditions. This may be attributed to the larger size of the flower in our studies, but the mechanism behind and consequence of the secondary peak is unknown. This implies that the delay model is a simplification of a mechanism with additional features.

**Figure 6.**
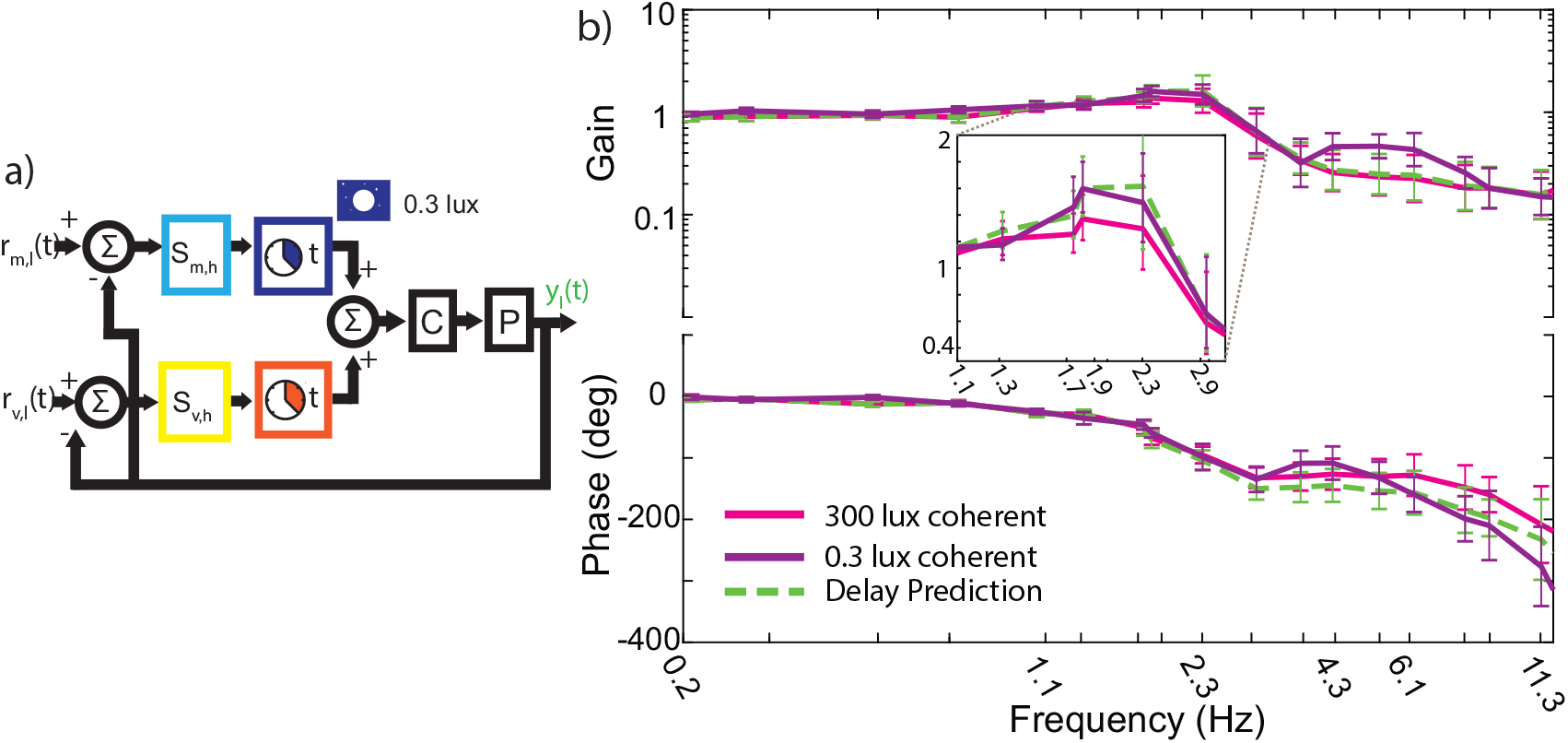
(a) Control diagram models for pure delays. Pure delay of 10ms is incompatible with multisensory data (Figure 3, Figure 4), while frequency-dependent gain-modulation (*K*_*m*_ and *K*_*v*_ in Figure 4) is consistent (b) Bode plots for high-luminance (300 lux, pink), low-lumiance (0.3 lux, purple) and delay-prediction (green dashed) transfer functions. Inset has linear y-axis, showing gain between 1.1Hz and 3.2Hz

While a delay in the coherent response is consistent with these features of coherent motion responses across the luminance levels, this does not mean that the visual and mechanosensory systems are purely delayed an equal amount. That would be incompatible with the observed shift in gains of the open-loop responses (Figure 6a). Instead, a frequency-dependent gain adjustment in both vision and mechanosensation can account for the change (in Figure 4, *K*_*m*_ and *K*_*v*_ are frequency-dependent gain knob). A more general model consisting of both variable-gain and delay tuning of both systems, may give comparable results. The pure-delay motif for coherent flower motion found in ***Sponberg et al. (2015***) results from a linear combination of gain and phase tuning of both open-loop, mechanosensory and visual sensorimotor branches. Specifically, the delay resulting from the visual system’s spatio-temporal summation at low luminance complemented by frequency-dependent gain and phase adjustments in the mechanosensory systems. This is reminiscent of ***Stöckl et al. (2017***), where gain and delay tuning was found to account for inter-species differences. Therefore, both within-species differences (this study) across contexts, and between-species differences (***Stöckl et al. (2017***)), are consistent with similar types of control adjustments.

## Discussion

### Multisensory Integration is Context Dependent for *Manduca sexta* Flower Tracking

We found that *Manduca sexta’s* proboscis-mechanosensory flower-tracking response is luminance-dependent, adapting in a manner separate from changes to vision. The mechanosensory and visual systems continue to combine linearly across light levels, and their coherently-combined effect is equivalent to a delay in the moth’s response. These luminance-dependent adaptations result in comparable flower-tracking performance across contexts. Thus, a linear combination of cues, individually re-weighted depending on luminance context, enables coherent tracking of flower motion in *Manduca sexta*. This multisensory integration and re-weighting of cues may be important because *Manduca sexta* experiences luminance changes across this range during foraging. The difference in peak-response frequencies of the mechanosensory and visual systems, combined with their frequency overlap, allows for the adaptive nature of the context-dependent response. Degradation of one system, such as increases in visual delay and loss of resolution with luminance decrease, can be partially accommodated by tuning the complementary (mechanosensory) system gain, at the appropriate speeds.

Multisensory integration may not be essential for flower-tracking in all species. Indeed, the diurnal hawkmoth, *Macroglossum stellatarum* relies mostly on its visual sense, and very little on proboscis cues, in response to looming flowers (***Farina et al. (1994***)), a fact underlined by its region-specific optical specializations (***Farina et al. (1995***), ***Warrant et al. (1999***)). In *Manduca sexta*, its longer proboscis may make for a more reliable organ for sensing (***Stöckl et al. (2017***)), which is especially desirable in low or variable light conditions. However, even *M. sexta* shows redundancy between the two systems (***Roth et al. (2016***)). This means that either system can, on its own, successfully produce flower-tracking. Having two redundant systems allows for compensation when visual salience is affected, such as when flying into shade or between evenings and late nights. The conditions which may lead to decreased mechanosensory salience are less well understood.

*Manduca sexta* feed from flowers moving in the wind, performing an energetically expensive and challenging hovering behavior. The energetic gains from successful feeding, however, have been estimated to more than make up for this loss (***Sprayberry and Daniel (2007***),***Stöckl and Kelber (2019***)), and are known to have fitness benefits in terms of the number of eggs laid (***Levin et al. (2016***), ***von Arx et al. (2013***)). In order to feed uninterrupted, they would benefit from tracking the frequencies of natural flower motion ***Sponberg et al. (2015***). The two sensorimotor systems used for flower tracking have appreciable luminance-dependent response-gain modulation at these frequencies (Figure 4). Hence the integration of visual and mechanosensory cues has an essential impact on the successful tracking of moving flowers.

The multisensory aspect of flower-tracking may be related to the use of the mechanosensory and visual systems in associated hawkmoth behaviors, such as proboscis placement (***Goyret and Kelber (2012***),***Goyret (2010***)), flower handling and learning (***Goyret and Raguso (2006***),***Deora et al. (2021***)). Switching between these behaviors seems to be gated by olfaction and other multimodal cues (***Raguso and Willis (2002***)). We find that luminance acts as an additional environmental cue, controlling the method of execution of a behavior, by re-weighting of sensorimotor gains. The re-weighting of sensory systems has been observed previously, in refuge-keeping of electric knifefish. Depending on the salience of electrosensory (conductance) and visual (light-level) information, ***Sutton et al. (2016***) found that the gain ratio of their two systems was modulated, with the information being combined in a roughly linear manner. While this study was done only at distinct probe frequencies, and not across the dynamic range, the re-weighting shows us that the same effective dynamics may emerge across distantly related taxa, when engaging in a similar behavior. This raises the question, is there a performance benefit to the linear combination of multisensory information?

### Sensorimotor Timescales of Visual and Mechanosensory Modalities Shift Inversely

Taken together our results show not only the visual tracking of object, but also the mechanosensing of objects changes with the background luminance. Each of the visual and mechanosensory responses show frequency tuning across overlapping frequency ranges (Figure 4). The width of the frequency bands over which each system’s gain is larger than the other’s, tells us the relative speed-tuning of the mechanosensory and visual systems. We see that the proboscis mechanosensory system operates over a slower timescale than the compound-eye visual system, with a gain crossover point defining each system’s bandwidth. As luminance increases, this crossover point moves to lower speeds. Thus we say that the visual system’s bandwidth increases, that of the mechanosensory system decreases. In other words, the visual system slows down (as evidenced by its peak shifting) enough to compensate for the loss of mechanosensory gain, at high luminance. The two systems continue to fulfill their roles of high- and low-speed response, respectively, but with their frequency bandwidths rescaled accordingly.

Bandwidth separation and complementary action are usually seen when two sensory systems have very different frequency tuning. Difference in timescales of visual (slow) and gyroscopic haltere (fast) responses in dipterans, enable optomotor reflexes and successful object-fixation behavior (***Mureli and Fox (2015***)). The optomotor response range of *drosophila* is expanded by the use of these two, differently tuned systems (***Sherman and Dickinson (2003***)). The use of high-speed mechanosensors also allows *drosophila* to stabilize their visually-controlled flight speed (***Fuller et al. (2014***)). In *Manduca sexta*, the function of high-frequency gyroscopic mechanosensation is performed by the antennae (***Sane et al. (2007***)). Similarly, in the closely-related species, Macroglossum stellatarum, ***Dahake et al. (2018***) noted the separation of antennae-guided stabilization, and visually-guided flower-tracking roles. In their study, the visual system could not compensate for the loss of antennae, even in bright lighting conditions. This led to the hypotheses of either spatial separation of the two systems, via parallel and independent descending pathways, or temporal separation (multiplexing) due to a difference in response speeds. Thus, they found that antennal mechanosensation is separate in its frequency response from visual and proboscis mediated cues. We find that the proboscis and eyes compensate for each other, and work in combination across contexts. This is possible, and even necessary due to the overlap in frequency bands between the eyes and proboscis. The shift in frequency bands in our system may be investigated further through electrophysiology in the proboscis and descending neurons

The proboscis and eyes, unlike the halteres-eyes and antennae-eyes systems, have considerable frequency-overlap, and fulfill similar roles. A closer analogy would be the widefield motion (optomotor) and flower-tracking systems, in the related hover-feeding species, *M. stellatarum*. When *M. stellatarum* feeds from a looming flower (***Farina et al. (1995***)), distance-stabilization (flower distance) and drift-compensation (widefield motion) responses have very different bandpass shapes, while sharing a common range of frequencies. Similarly, the response to rotating widefield motion (***Kern (1998***),***Kern and Varju (1998***)) is strongly peaked. These behaviors may be in conflict or coherent, depending on the context of motion (intent) of the moth and changes in its environment. Thus, even more extreme rescaling of sensory information has been observed here, dependent on internal and external context (albeit for the same sensory system, i.e. vision).

### Luminance dependent delays in visual processing can arise form multisensory reweighting

Luminance-dependent changes in the visual system have been studied extensively, and at various stages, in insects. Matched filtering (***Kohn et al. (2018***)) has been found to result in speed tuning of sensory systems depending on their natural ecology, which may in turn span different luminance contexts. Light-level affects the temporal frequency tuning of lobula plate tangential cells (LPTCs) (***O’Carroll and Warrant (2011***)) of *Deilephila elpenor*, and summation of spatial information by lamina monopolar cells (LMCs) (***Stöckl et al. (2020***)), in addition to physical dark-adaptation in the eyes. The result of such adaptation, along with possible differences in wing kinematics and control, can result as an effective delay in flower tracking for *M. sexta* (***O’Carroll and Warrant (2011***); ***Sponberg et al. (2015***)).

Our observation, that coherent motion tracking is consistent with previously characterized delay (***Sponberg et al. (2015***)), gives us a model for how the visual and mechanical systems must combine: the result of their respective luminance-tuning adds to give an overall system delay. Flowertracking behavior has been previously studied in the linear regime for three species of hawkmoths (***Stöckl et al. (2017***)), and inter-species differences are matched to natural ecology, with speciesdependent gain scaling and luminance-tuned delays. We found that gain and delay tuning of sub-systems, can account for luminance dependent tracking differences within a species as well. Electrophysiological studies of sensory integration, combined with analyses of flight EMGs and dynamics, may shed light on how this template is achieved.

We found that the delay is the result of multisensory integration and control, and the proboscis mechanosensory system has appreciable luminance-dependent gain-modulation. The corresponding changes in the proboscis, if any, have yet to be studied. Alternatively, an inverse set of changes closer to flight control centers, could also account for this gain tuning, without direct modulation of proboscis sensors. If true, this could be a general locomotion strategy used by animals, whereby multiple sensory systems can robustly combine to control different speeds of action, through changes in a subset of the sensors, and corresponding changes in output control sites.

## Methods and Materials

### Robotic flower for sensory conflict experiments

Following the methods described by ***Roth et al. (2016***), we used a two-part robotic flower to generate coherent motion as well as sensory conflict stimuli. The motion of the flower facade provided visual stimuli, and the nectary motion provided mechanosensory cues. The robotic flower’s dimensions matched those used by ***Roth et al. (2016***). The facade and nectary were 3d printed out of ABS P430 using a Stratasys® uPrint® 3D printer, and were mounted on 16 cm carbon fiber rods. The flower facade had a slit wide enough to not interfere with proboscis motion (2mm). The rubbing of the facade against the proboscis would suggest a phase-leading derivative (velocity) response, where as we observe, low phase lags consistent with a proportional (positional) response. Thus, movement of the facade did not deflect the stationary proboscis in the absence of nectary movement. Two stepper motors (57sth56-2804mb, Dongzheng. Motor Co. LTD; China) were used to provide independent movement stimuli. The nectary was painted black to minimize visual distraction. Our robotic flower differed from the ones used by ***Sponberg et al. (2015***), in its diameter (9.5cm vs 4.5cm). The larger diameter of our robotic flower was necessary to generate the sensory conflict stimuli. Hence, the applicability of the control-system delay theory to our paradigm would require further evaluation.

### Sum-of-sines Stimuli

Sum of sines stimuli were provided, with 20 frequency components (being prime multiples of 0.1Hz), logarithmically spaced, from 0.2Hz to 19.9Hz. This stimulus was presumed to be pseudorandom, so that it could not be predicted by the moth. Stimulus amplitudes were chosen such that velocity amplitudes were equal at each frequency. This velocity was similar to the one used by ***Sponberg et al. (2015***). Phase difference for each component sine waveform was randomized, and kept constant across trials for a given luminance level. We tested 3 stimulus conditions: Co-herent motion, M-moving and V-moving. In coherent motion, both the facade and nectary moved together. In the M-moving case, the facade was stationary, while the nectary was moved as a sum of sines. Conversely, in the V-moving case, the facade was moved while the nectary remained stationary.

### Animal Handling and Video Recording

We obtained *Manduca sexta* pupae from the University of Washington. Within 2 to 3 days posteclosion, 8 moths were placed successively in a flight chamber, with the robotic flower facade being illuminated at 300lux. The light source used was the LED light, Neewer CN-126, with 2 attached neutral density filter layers. For ease of tracking, we painted a white spot (Prang White Paint, Dixon Ticonderoga Co.) on each moth’s thorax before the start of every trial. The MATLAB plug-in, DLT-DV was used to track the position of the thorax of each individual. We painted a black spot on top of the flower facade, and a white spot on top of the nectary (out of view of the feeding moth) for automated tracking. Tracks were validated manually. Movement stimuli were set to ramp up linearly in amplitude, over 2 seconds during the beginning and end of recordings, to prevent sudden impulsive stimuli and motor saturation. Sum of sines stimuli were provided for 20 seconds, in order to get four repeats of the lowest frequency. A Fastec camera was used to record the trials (top view), at 2000fps. A seven-component scent (***Campos et al. (2015***) 0.9-1.7uL) was placed on a Q-tip™ to increase the likelihood of feeding behavior. Stimulus order was randomized, and each individual was made to fly once every day, over 3-4 days. Hence, each feeding individual was presented with all 3 stimuli. Individuals that did not feed on at least 3 of these days were excluded from analysis. Individuals were weighted before and after each trial, and the decrease in nectary weight was compared with increase in moth weight, while the nectary was refilled for each trial. The procedure was repeated for 10 more feeding individuals. Out of 18 trials, 12 were identified to coherently track and feed from flowers, and did not suffer excessive visible wing-damage, and these were included in our analysis.

Low-luminance (0.3 lux) tracking results were taken directly from Roth et al’s dataset.

### Linear combination of cues in Individuals

We collected triads of data at 300 lux, with one trial for each stimulus condition per individual. Therefore, we were able to evaluate the linear combination of cues for each individual moth. The predicted coherent response is the sum of M-moving and V-moving responses:

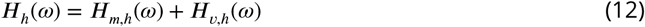

We find this to be the case for every individual tested

### Frequency-Domain Analysis and Statistics

#### Transfer Functions

We used the same statistical analysis technique as ***Roth et al. (2016***) for complex-valued transfer-functions. Briefly, the transfer function H(*ω*) is given by:

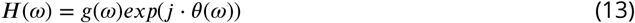

The log-space representation of this transfer-function is:

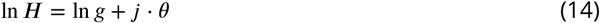

In log-space, we calculate the mean and standard deviation of the gain, and then return to the untransformed complex plane:

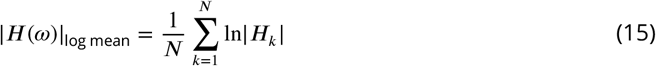

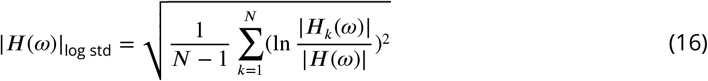

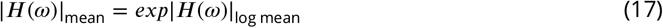

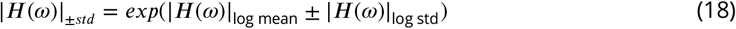

The phase of the transfer functions is given by:

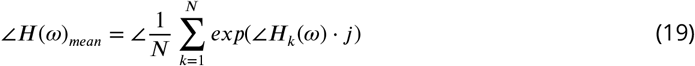

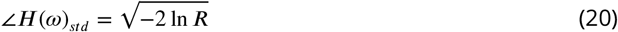

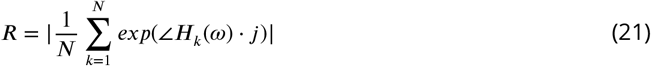

Statistics on predicted values of transfer functions are done by propagation of uncertainty analysis, similar to ***Roth et al. (2016***). These include predictions for the linear sum, the open-loop transfer functions, and the delay predictions.

For low luminance, the data was unpaired. Therefore, to calculate the open-loop frequency response functions, *G*_*m,l*_ and *G*_*v,l*_, we made every possible combination of the 8×8×8 individuals (8 experimental trials each for M, V and MV stimuli). This gave us 512 “paired” triads of data. We then extended equations (6) and (7) with these 512 rows of triads, and estimated the open-loop functions by solving this version of equation (7). These synthetic transfer functions provide an upper bound on the variance.

#### Time Domain and Fourier Amplitude

The time-series of individuals are aligned to the start of stimuli, and a timepoint-specific standard deviation is found for each stimulus condition (Figure 1b).

The Fourier amplitudes for these timeseries have peaks at driving frequencies. Peak significance is estimated similar to ***Sponberg et al. (2015***), assuming white-noise as null signal. The frequency range for significant peaks depends on the individual and the stimulus condition (coherent or conflict). Therefore, only frequencies up to 6.1 Hz are used in analyses, since these frequencies have significant Fourier peaks.

## Acknowledgements

We would like to thank Eatai Roth and Nowan Cowan for helpful comments and suggestions. Funding was provided by Air Force Office of Scientific Research grant FA9550-22-1-0315, NSF Faculty Early Career Development Award (Award no. 1554790), and NSF Physics of Living Systems SAVI student research network (GT node grant no. 1205878).

