## Supplementary Information for "Context dependent multisensory integration: Mechanosensation depends on luminance for robust performance"

### Supplementary Materials

**Individual Linear Summation**

In our dataset of high luminance (300 lux) flower-tracking, each of the 12 moths performed hover-feeding in all 3 conditions: Mechanical motion, Visual motion, and Coupled motion. Therefore, we could verify linear summation of the frequency response functions for every individual moth. Over the range of natural frequencies, the coupled motion response matched the linear sum prediction for most individuals.

Figure S1 (i-xii) illustrates this for Moths 1 to 12.

S1 (i)

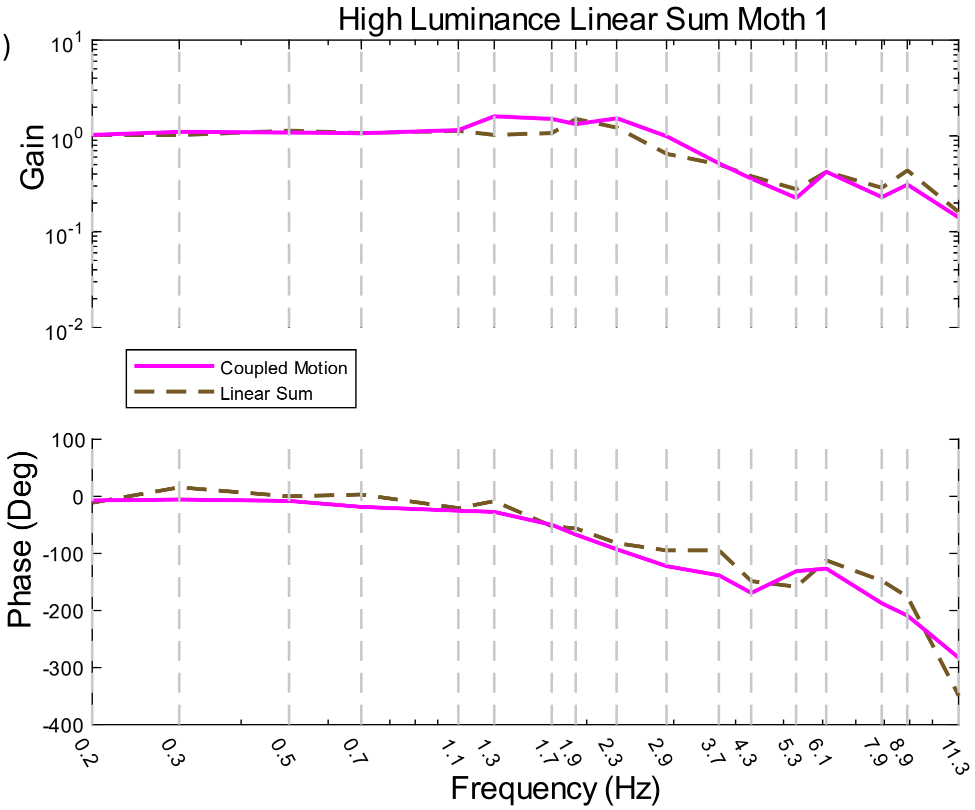

S1 (ii)

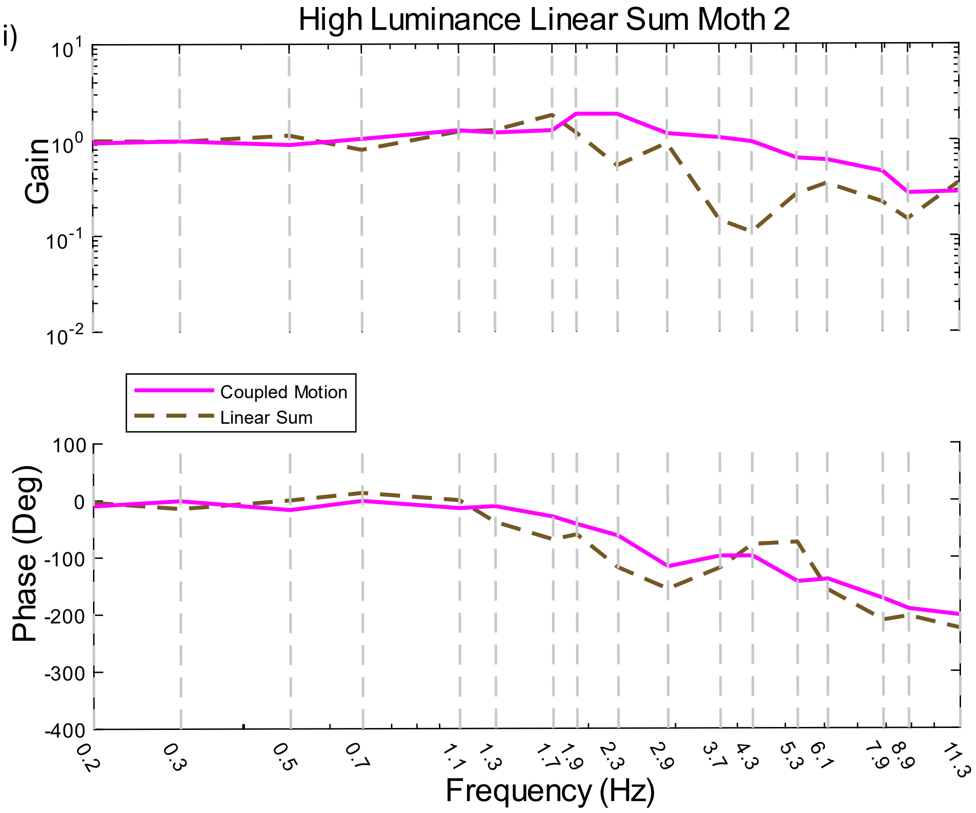
High Luminance Linear Sum Moth 2
Gain
Phase (Deg)
Frequency (Hz)
Coupled Motion
Linear Sum
0.2
0.3
0.5
0.7
1.1
1.3
1.7
1.9
2.3
2.9
3.7
4.3
5.3
6.1
7.9
8.9
11.3

S1 (iii)

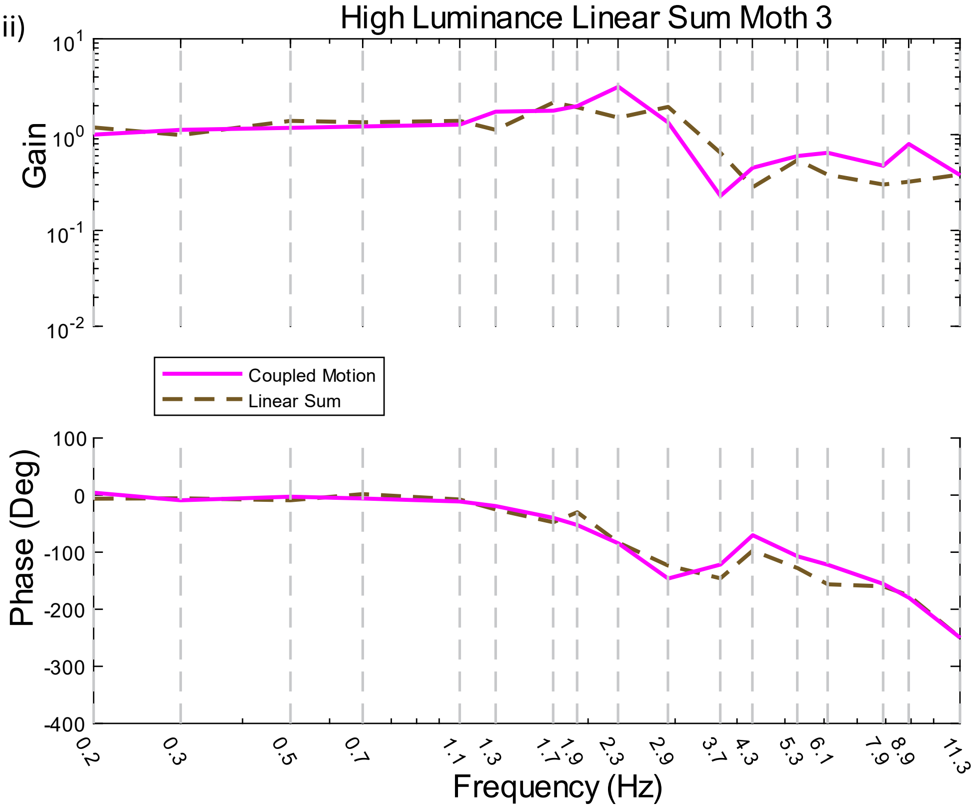
High Luminance Linear Sum Moth 3
Gain
Phase (Deg)
Frequency (Hz)
Coupled Motion
Linear Sum
0.2
0.3
0.5
0.7
1.1
1.3
1.7
1.9
2.3
2.9
3.7
4.3
5.3
6.1
7.9
8.9
11.3

S1 (iv)

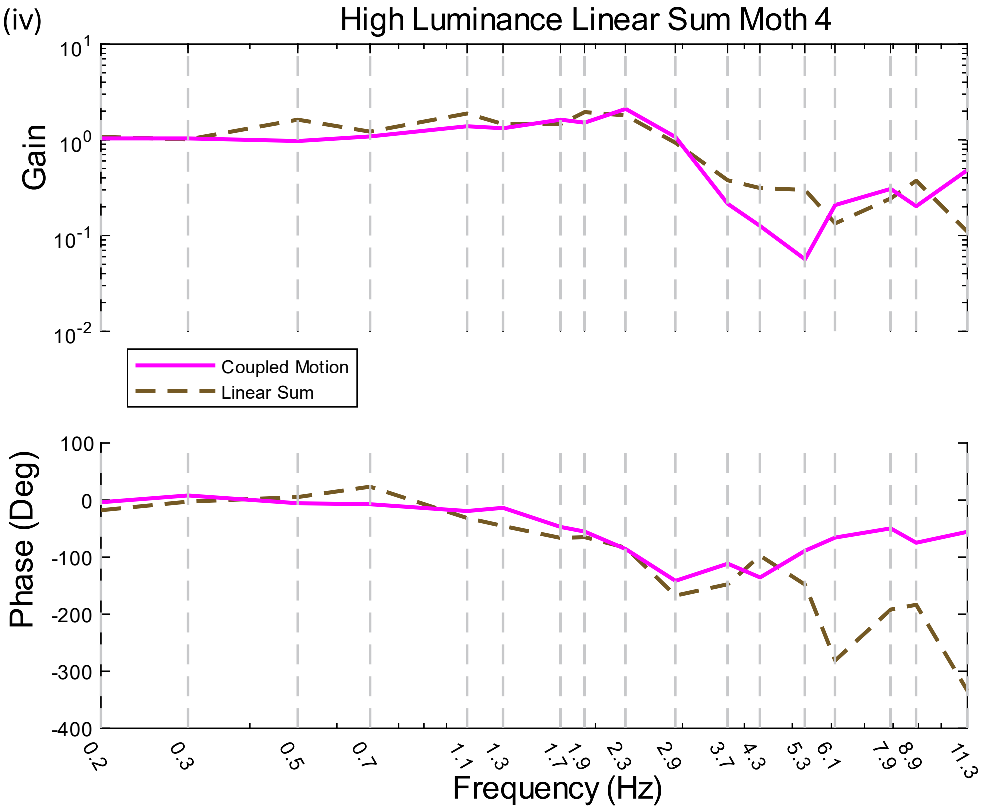
High Luminance Linear Sum Moth 4
Gain
Coupled Motion
Linear Sum
Phase (Deg)
Frequency (Hz)

S1 (v)

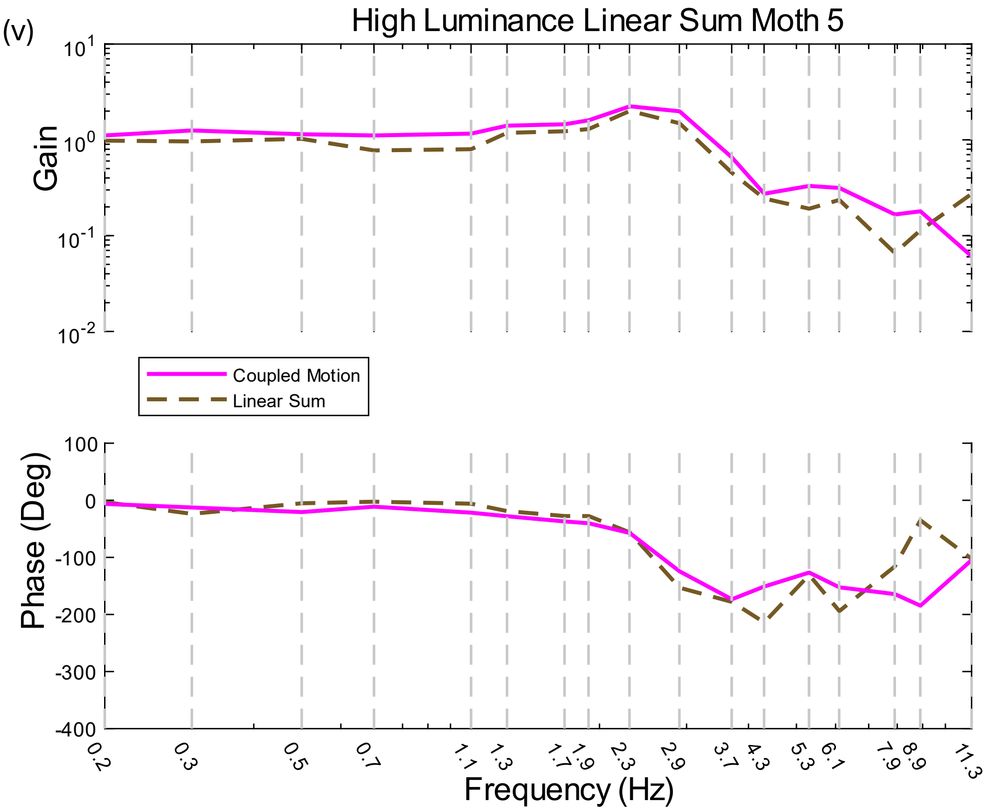
High Luminance Linear Sum Moth 5
Gain
Coupled Motion
Linear Sum
Phase (Deg)
Frequency (Hz)

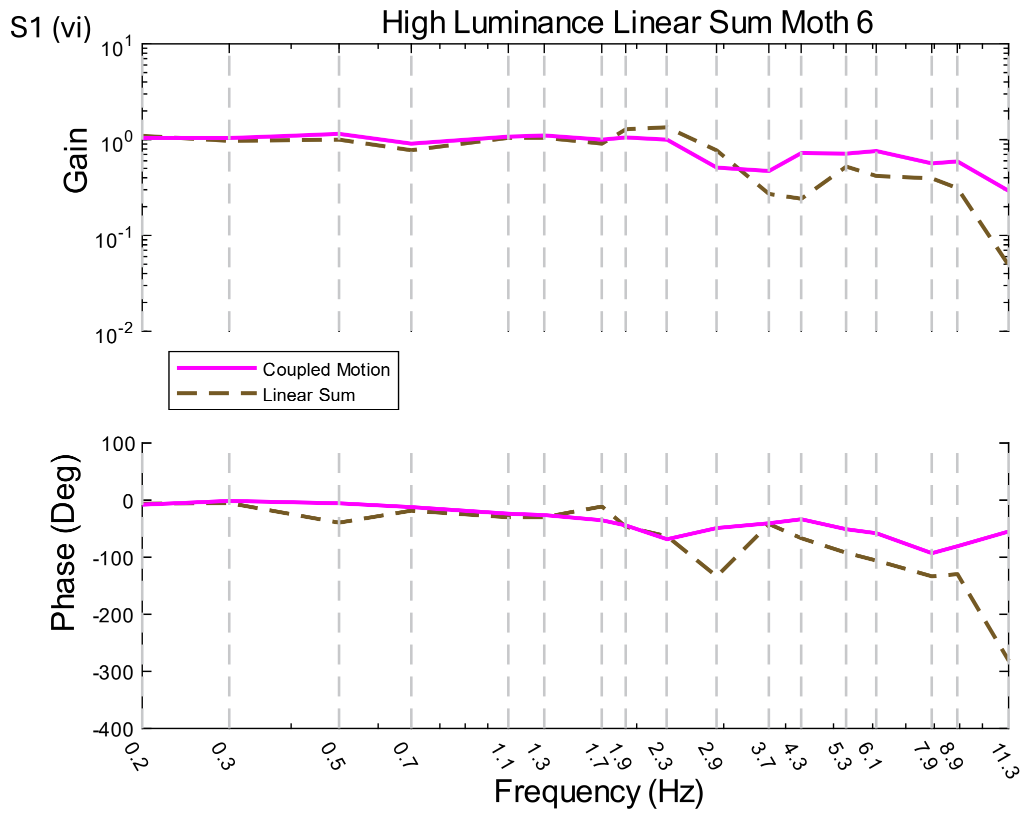
S1 (vi)
High Luminance Linear Sum Moth 6
Gain
Coupled Motion
Linear Sum
Phase (Deg)
Frequency (Hz)

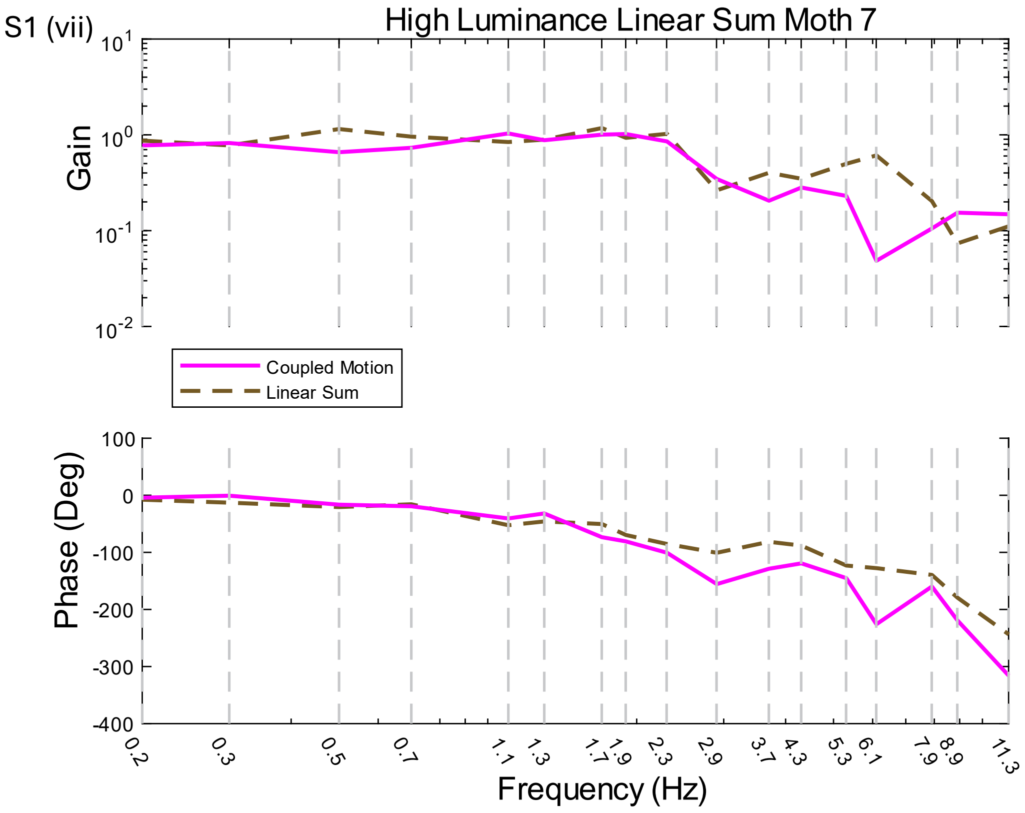
S1 (vii)
High Luminance Linear Sum Moth 7
Gain
Coupled Motion
Linear Sum
Phase (Deg)
Frequency (Hz)

S1 (viii)

High Luminance Linear Sum Moth 8

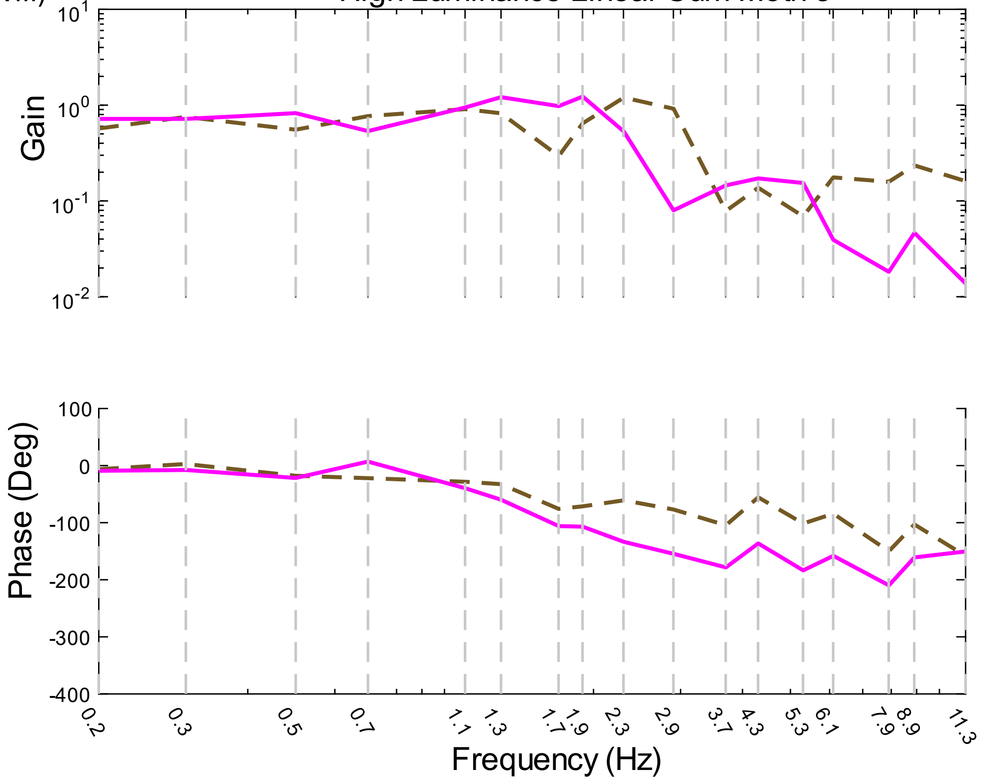


S1 (ix)

High Luminance Linear Sum Moth 9

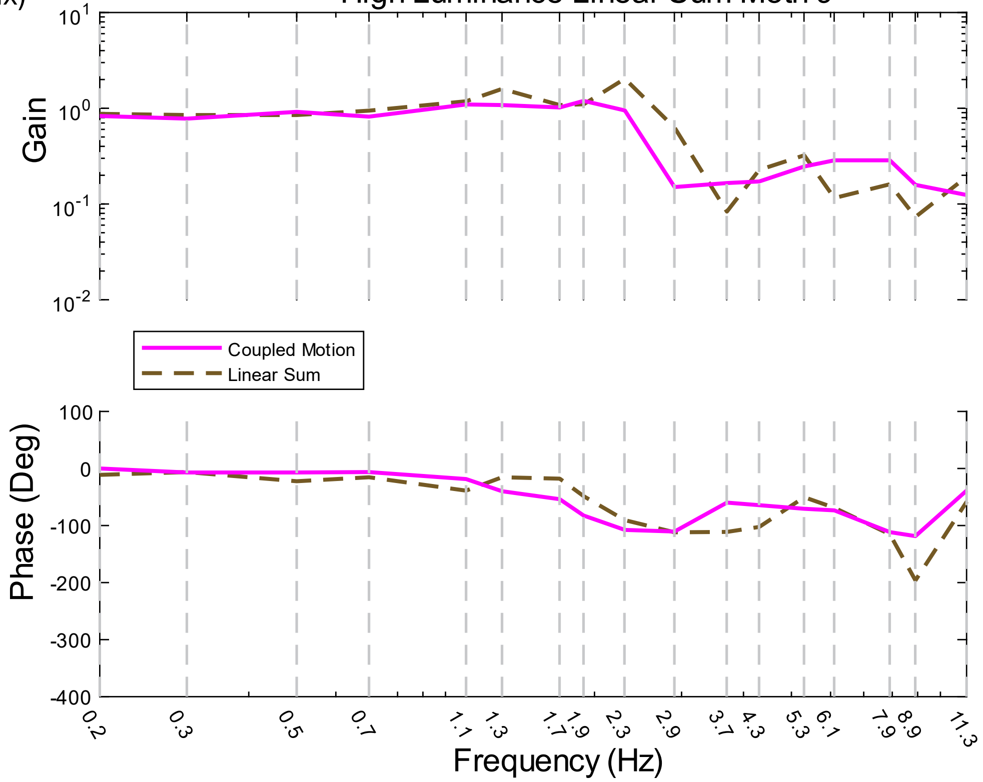

S1 (x)

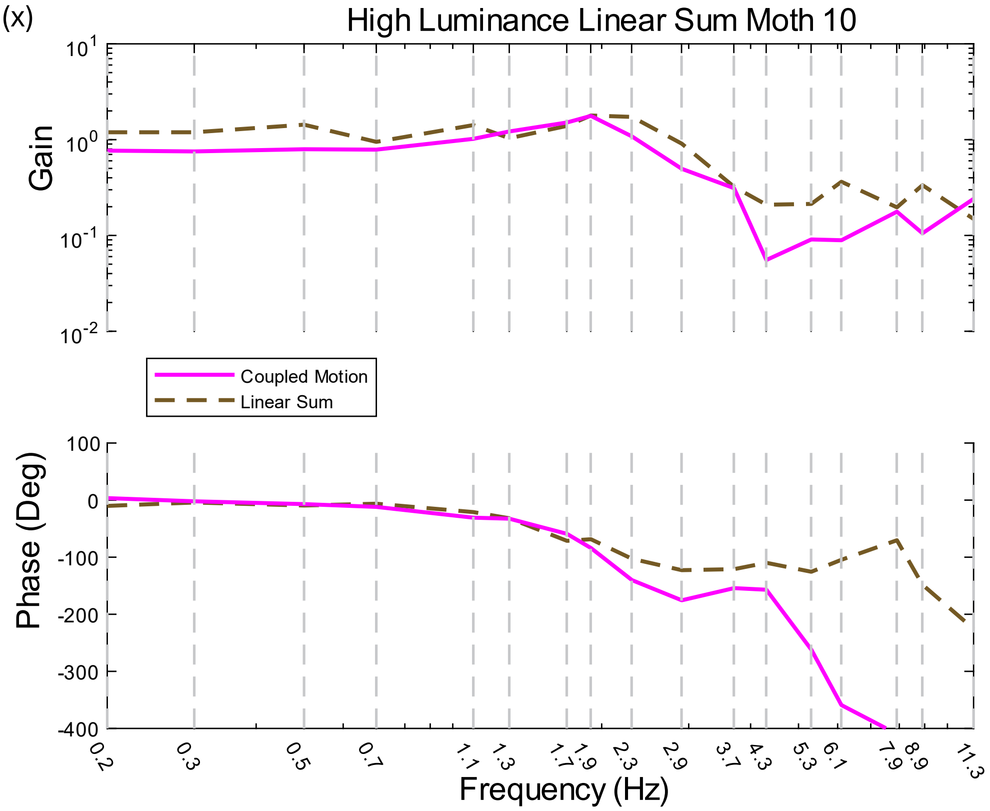

High Luminance Linear Sum Moth 10
Gain
Coupled Motion
Linear Sum
Phase (Deg)
Frequency (Hz)


S1 (xi)

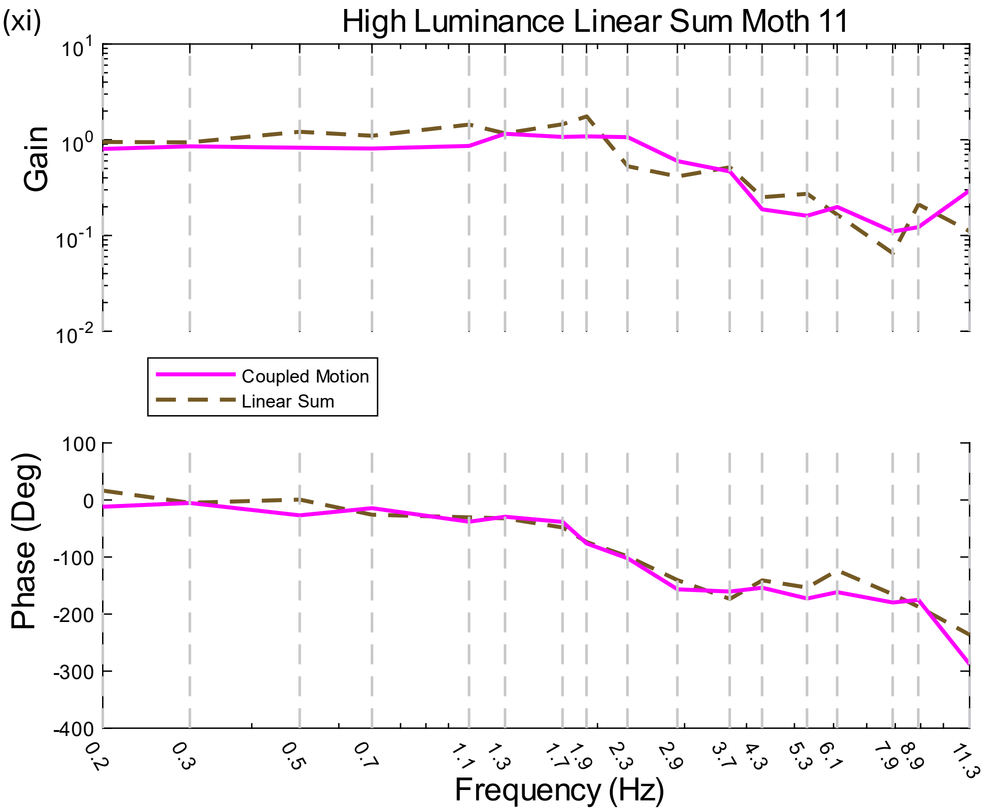

High Luminance Linear Sum Moth 11
Gain
Coupled Motion
Linear Sum
Phase (Deg)
Frequency (Hz)

S1 (xii)

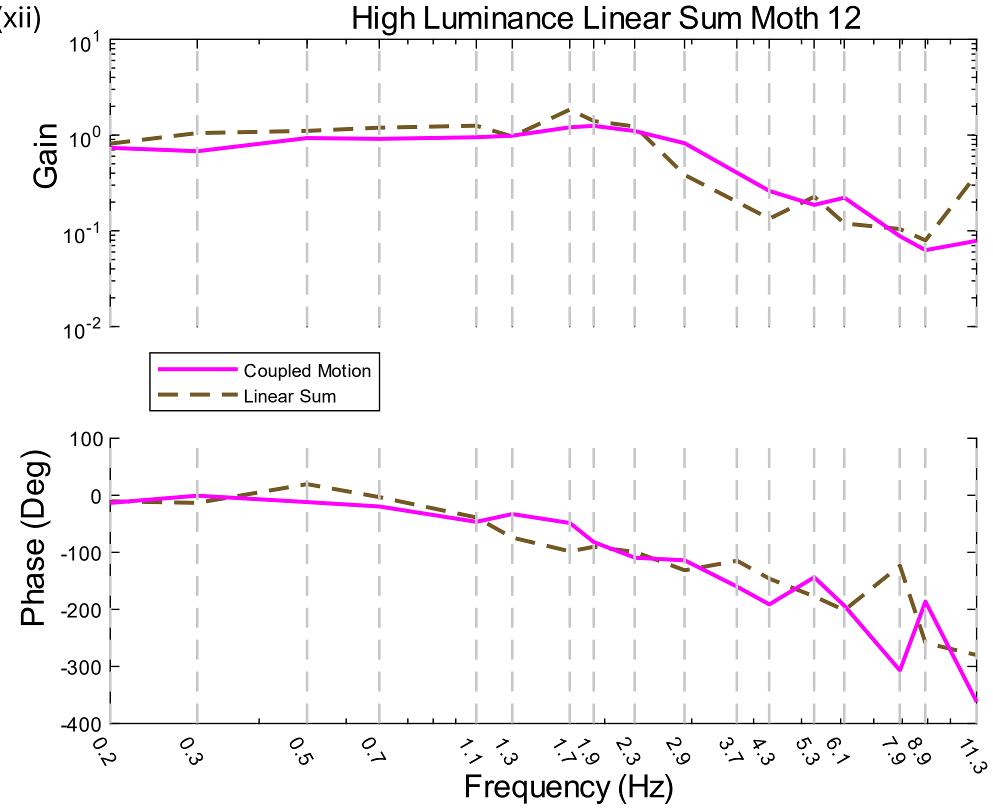


**Figure S1 (i-xii) Individual Linear Sum:** At 300 lux conditions, all 12 moths show good agreement between coupled motion frequency response and linear sum prediction, over the range of natural frequencies.

**Statistics of gain and phase**

The transfer function

$$H(\omega) = \exp\left(g(\omega) + j \cdot \theta(\omega)\right)$$

We take its log-space representation, as seen in the Bode plots, for statistics

<u>Gain Statistics</u>

The gain, as a function of frequency, has log-mean:

$$|H(\omega)|_{logmean} = \frac{1}{N} \cdot \sum_{k=1}^{N} \log(|H_k(\omega)|)$$

where k is moth-number, from 1 to 12 for 300 lux data, and from 1 to 8 for 0.3 lux data

Gain mean is given by

$$|H(\omega)|_{mean} = \exp\left(|H(\omega)_{logmean}|\right)$$

The log standard deviation of the mean is

$$|H(\omega)|_{logstd} = \sqrt{\frac{1}{N-1} \cdot \sum_{k=1}^{N} \left( \log \left( \frac{|H_k(\omega)|}{|H(\omega)_{mean}|} \right) \right)^2}$$

and the mean gain with standard deviation is

$$|H(\omega)|_{\pm std} = \exp\left(|\mathrm{H}(\omega)|_{\mathrm{logmean}} \pm |\mathrm{H}(\omega)|_{\mathrm{logstd}}\right)$$

<u>Phase statistics</u>

Following Roth et al, we calculate the mean phase as

$$\angle H(\omega)_{mean} = \angle \frac{1}{N} \cdot \sum_{k=1}^{N} \exp\left(j \cdot \angle H_k(\omega)\right)$$

and the standard deviation as

$$\angle H(\omega)_{std} = \sqrt{-2 \cdot \log(R)}$$

with vector strength

$$R = \left| \frac{1}{N} \cdot \sum_{k=1}^{N} \exp\left(j \cdot \angle H_k(\omega)\right) \right|$$

<u>Linear Sum Prediction Statistics</u>

The mechanosensory and visual gain and standard deviation of gain, in log representation, is given by

$$\bar{g}_m(\omega) = \frac{1}{N} \cdot \sum_{k=1}^{N} \log\left(\left|H_{m,k}(\omega)\right|\right)$$

$$\bar{\sigma}_{gain,m}(\omega) = \sqrt{\frac{1}{N-1} \cdot \sum_{k=1}^{N} \left( \log \left( \frac{|H_{m,k}(\omega)|}{\exp\left(\bar{g}_m(\omega)\right)} \right) \right)^2}$$

and

$$\bar{g}_v(\omega) = \frac{1}{N} \cdot \sum_{k=1}^{N} \log\left(\left|H_{v,k}(\omega)\right|\right)$$

$$\bar{\sigma}_{gain,v}(\omega) = \sqrt{\frac{1}{N-1} \cdot \sum_{k=1}^{N} \left( \log \left( \frac{|H_{v,k}(\omega)|}{\exp\left(\bar{g}_v(\omega)\right)} \right) \right)^2}$$

The mechanosensory and visual phase and standard deviation of phase, in log representation, is given by

$$\bar{\theta}_m = \angle \left( \frac{1}{N} \cdot \sum_{k=1}^{N} \exp\left(j \cdot \angle H_{m,k}(\omega)\right) \right)$$

$$\bar{\sigma}_{phase,m} = \sqrt{-2 \cdot \log \left(\frac{1}{N} \cdot \sum_{k=1}^{N} \exp\left(j \cdot \angle H_{m,k}(\omega)\right)\right)}$$

and

$$\bar{\theta}_v = \angle\left(\frac{1}{N} \cdot \sum_{k=1}^{N} \exp\left(j \cdot \angle H_{v,k}(\omega)\right)\right)$$

$$\bar{\sigma}_{phase,v} = \sqrt{-2 \cdot \log \left(\frac{1}{N} \cdot \sum_{k=1}^{N} \exp\left(j \cdot \angle H_{v,k}(\omega)\right)\right)}$$

The linear sum prediction states that mechanosensory and visual frequency response functions add to give coherent motion response

$$H_m(\omega) + H_v(\omega) = \widehat{H}(\omega)$$

$$\widehat{H}(\omega) = \exp\left(\hat{g}(\omega) + j \cdot \hat{\theta}(\omega)\right)$$

Therefore, the predicted gain and phase are

$$\hat{g}(\omega) = \log\left(\left|\exp(\bar{g}_m + j \cdot \bar{\theta}_m) + \exp(\bar{g}_v + j \cdot \bar{\theta}_v)\right|\right)$$
$$= \log\left(\sqrt{\Big(\big((\exp(\bar{g}_m) \cdot \cos(\bar{\theta}_m)) + (\exp(\bar{g}_v) \cdot \cos(\bar{\theta}_v))\Big)^2 + \Big((\exp(\bar{g}_m) \cdot \sin(\bar{\theta}_m)) + (\exp(\bar{g}_v) \cdot \sin(\bar{\theta}_v))\Big)^2}\right)$$

$$\hat{\theta}(\omega) = \angle\left((\exp(\bar{g}_m + j \cdot \bar{\theta}_m) + \exp(\bar{g}_v + j \cdot \bar{\theta}_v)\right)$$
$$= \arctan\left(\frac{(\exp(\bar{g}_m) \cdot \sin(\bar{\theta}_m)) + (\exp(\bar{g}_v) \cdot \sin(\bar{\theta}_v))}{((\exp(\bar{g}_m) \cdot \cos(\bar{\theta}_m)) + (\exp(\bar{g}_v) \cdot \cos(\bar{\theta}_v))}\right)$$

For the standard deviation of the predicted linear-sum response, we use propagation-of-uncertainty analysis.

This uses the Jacobian evaluations for gain and phase

$$J_{\hat{g},\bar{g}_m} = \frac{\partial \hat{g}}{\partial \bar{g}_m}, \quad J_{\hat{\theta},\bar{g}_m} = \frac{\partial \hat{\theta}}{\partial \bar{g}_m}$$

$$J_{\hat{g},\bar{\theta}_m} = \frac{\partial \hat{g}}{\partial \bar{\theta}_m}, \quad J_{\hat{\theta},\bar{\theta}_m} = \frac{\partial \hat{\theta}}{\partial \bar{\theta}_m}$$

$$J_{\hat{g},\bar{g}_v} = \frac{\partial \hat{g}}{\partial \bar{g}_v}, \quad J_{\hat{\theta},\bar{g}_v} = \frac{\partial \hat{\theta}}{\partial \bar{g}_v}$$

$$J_{\hat{g},\bar{\theta}_v} = \frac{\partial \hat{g}}{\partial \bar{\theta}_v}, \quad J_{\hat{\theta},\bar{\theta}_v} = \frac{\partial \hat{\theta}}{\partial \bar{\theta}_v}$$

We define,

$$J_g = \left[J_{\hat{g},\bar{g}_m}, \quad J_{\hat{g},\bar{\theta}_m}, \quad J_{\hat{g},\bar{g}_v}, \quad J_{\hat{g},\bar{\theta}_v}\right]$$

and

$$J_\theta = \left[J_{\hat{\theta},\bar{g}_m}, \quad J_{\hat{\theta},\bar{\theta}_m}, \quad J_{\hat{\theta},\bar{g}_v}, \quad J_{\hat{\theta},\bar{\theta}_v}\right]$$

Using the measured mechanosensory and visual standard deviations, we define the diagonal matrix

$$\Sigma = \begin{pmatrix} \bar{\sigma}_{gain,m} & 0 & 0 & 0 \\ 0 & \bar{\sigma}_{phase,m} & 0 & 0 \\ 0 & 0 & \bar{\sigma}_{gain,v} & 0 \\ 0 & 0 & 0 & \bar{\sigma}_{phase,v} \end{pmatrix}$$

At each frequency, the resulting standard deviation for the linear sum prediction is

$$\hat{\sigma}_{gain,linearsum} = \sqrt{J_g \Sigma^2 J_g^T}$$

$$\hat{\sigma}_{phase,linearsum} = \sqrt{J_\theta \Sigma^2 J_\theta^T}$$

Open Loop Response Estimate for High Luminance

For the 300 lux (high luminance) condition, we have triplets of Nectary-Moving (M), Façade-Moving (V) and Coherent-Motion data for each of the 12 individuals. Therefore, we calculate the open-loop response for each individual moth.

$$\hat{G}_m = \frac{H_m}{1-H} = \exp\left(\hat{g}_{m,open} + j \cdot \hat{\theta}_{m,open}\right)$$

$$\hat{G}_v = \frac{H_v}{1-H} = \exp\left(\hat{g}_{v,open} + j \cdot \hat{\theta}_{v,open}\right)$$

Since

$$(1-H) \cdot \hat{G}_m + (1-H) \cdot \hat{G}_v = H$$

$$(1-H_m) \cdot \hat{G}_m + (-H_m) \cdot \hat{G}_v = H_m$$

$$(-H_v) \cdot \hat{G}_m + (1-H_v) \cdot \hat{G}_v = H_v$$

We define E, $\psi$ and $\hat{G}$

$$E(\omega)\hat{G}(\omega) = \psi(\omega)$$

$$E = \begin{pmatrix} 1-H & 1-H \\ 1-H_m & -H_m \\ -H_v & 1-H_v \end{pmatrix}$$

$$\hat{G} = \begin{pmatrix} \hat{G}_m \\ \hat{G}_v \end{pmatrix}$$

$$\psi = \begin{pmatrix} H \\ H_m \\ H_v \end{pmatrix}$$

$$\hat{G} = E \backslash \psi$$

Using MATLAB's *mldivide (\)*, we compute the least squares solution for $\hat{G}$, for each of the 12 moths. Then, we compute the gain and phase means and standard deviations in log-space as before.

Open Loop Response Estimate for Low Luminance

For the 0.3 lux (low luminance) condition, we do not have triplets of experiment conditions for each individual moth. In these experiments, 8 different individuals were used for each of the 3 conditions (M, V and coherent motion), for a total of 24.

To find an upper bound on the variability for this data, we formed artificial triplets from these 24 trials, with 1 individual from each of the 3 conditions. This results in 8 x 8 x 8 = 512 triplets.

Treating each of these 512 triplets like data from a single individual, we have for the $i^{th}$ triplet

$$E_i(\omega)\hat{G}_i(\omega) = \psi_i(\omega)$$

$$E_i = \begin{pmatrix} 1 - H_i & 1 - H_i \\ 1 - H_{m,i} & -H_{m,i} \\ -H_{v,i} & 1 - H_{v,i} \end{pmatrix}$$

$$\hat{G}_i = \begin{pmatrix} \hat{G}_{m,i} \\ \hat{G}_{v,i} \end{pmatrix}$$

$$\psi_i = \begin{pmatrix} H_i \\ H_{m,i} \\ H_{v,i} \end{pmatrix}$$

Then, as before, we calculate the least squares solution for $\hat{G}_i$ using MATLAB's *mldivide*, and estimate the mean and standard deviation for the gain and phase of $\hat{G}_m$ and $\hat{G}_v$ in the log-space, using N = 8 (and not 512) for standard-deviation calculations.
